# Differential Expression of N- and O-glycans on HeLa Cells-Revealed by Direct Fluorescent Glycan Labeling with Recombinant Sialyltransferases

**DOI:** 10.1101/580571

**Authors:** Zhengliang L Wu, Anthony D Person, Andrew J. Burton, Ravinder Singh, Barbara Burroughs, Dan Fryxell, Timothy J. Tatge, Timothy Manning, Guoping Wu, Karl A.D. Swift, Vassili Kalabokis

**Affiliations:** Bio-techne, R&D Systems, Inc. 614 McKinley Place N.E. Minneapolis, MN, 55413, USA; Bio-Techne, Tocris Bioscience, The Watkins Building, Atlantic Road, Avonmouth, Bristol, BS11 9QD, UK

**Keywords:** Glycan imaging, N-glycan, O-glycan, Sialyltransferase, T antigen

## Abstract

Cells are covered with glycans. The expression and distribution of specific glycans on cell surface are important for various cellular functions. Imaging these glycans is essential to elucidate their biological roles. Here, utilizing enzymatic incorporation of fluorophore-conjugated sialic acids, dubbed as <u>d</u>irect <u>f</u>luorescent <u>g</u>lycan <u>l</u>abeling (DFGL), we report the imaging of N- and O-glycans and particularly tumor specific sialyl T antigen on HeLa cells. It is found that while Core-1 O-glycans are relatively evenly distributed on cells, the expression of N-glycans tend to be more peripheral. More interestingly, the expression of sialyl T antigen displays random and sporadic patterns. In technique side, DFGL allows convenient labeling or imaging of various glycans on intact glycoproteins or cell surfaces with variety of fluorescent dyes.

## Introduction

The most common glycans are N-glycans that are linked to asparagine residues (Stanley, P., Taniguchi, N., et al. 2015) and O-Glycans that are linked to serine/threonine residues (Brockhausen, I. and Stanley, P. 2015). They are commonly found on cell surface, and involved in cell-cell, cell-extracellular matrix, and cell-pathogen interactions, and therefore play important roles in cell growth, differentiation and migration, innate immunity, and pathogenesis (Tabak, L.A. 2010, Taniguchi, N. and Kizuka, Y. 2015, Varki, A. 2017). When sialylated, they bind to numerous selectins expressed on mammalian cells (Crocker, P.R., Clark, E.A., et al. 1998) and function as binding targets for a large number of pathogenic organisms and their toxins (Schauer, R. 2009, Varki, A. 2008). They are also involved in cancer malignancy and inflammation, and therefore are potential targets for therapeutic and diagnostic intervention (Dube, D.H. and Bertozzi, C.R. 2005, Magalhaes, A., Duarte, H.O., et al. 2017). For their important biological functions and therapeutic and diagnostic applications, specific imaging or detection of these glycans is especially valuable (Chen, X. and Varki, A. 2010, Cummings, R.D. and Pierce, J.M. 2014).

N-glycans and O-glycans can be sialylated by multiple sialyltransferases (Harduin-Lepers, A., Mollicone, R., et al. 2005) (Fig. 1**A** and 1**B**). N-glycan sialylation typically occurs on Gal residues and is mediated by the N-glycan specific α-2,6-sialyltransferase 1 (ST6Gal1) (Weinstein, J., Lee, E.U., et al. 1987) and maybe α-2,3-sialyltransferase 4 (ST3Gal4) (Mereiter, S., Magalhaes, A., et al. 2016). O-glycan synthesis is initiated with an O-GalNAc residue that can be elongated to Core-1 O-glycan (T antigen, Galβ1,3-GalNAc-O-S/T) (Ju, T. and Cummings, R.D. 2005) that can be further sialylated by the O-glycan specific α-2,3-sialyltransferase 1 and 2 (ST3Gal1 and ST3Gal2) to generate sialyl-T antigen (sT antigen, Siaα2,3-Galβ1-3GalNAc-O-S/T) (Ju, T., Lanneau, G.S., et al. 2008, Kitagawa, H. and Paulson, J.C. 1994). O-GalNAc residue can also be directly sialylated at C-6 by a family of α-N-acetylgalactosaminide α-2,6-sialyltransferases (ST6GalNAc). ST6GalNAc4 is strictly active on sT antigen and responsible for disialylated T antigen expression (Harduin-Lepers, A., Stokes, D.C., et al. 2000).

**Figure 1.**
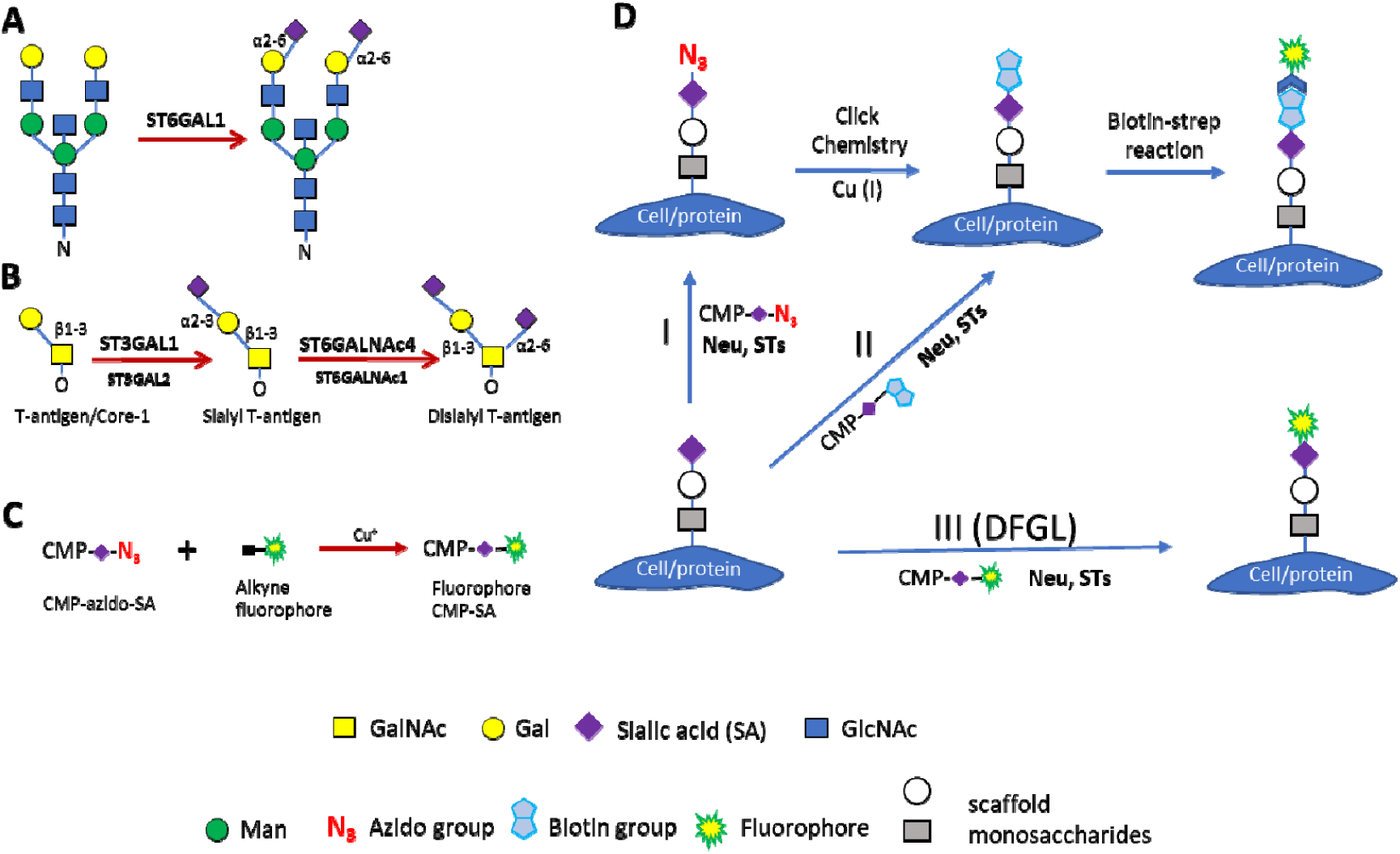
Sialylation reactions on N-glycans (A) and O-glycans (B) utilized in this report, method of fluorophore-conjugated CMP-Sialic acid preparation (C), and methods for glycan labeling (D). In **D,** three methods are depicted for glycan labeling and imaging. In method I, azido-sialic acid is incorporated into target glycans, followed by click chemistry reaction to attach a biotin molecule and then labeled by fluorophore-conjugated streptavidin. In method II, biotinylated sialic acid is incorporated into sialoglycans and then labeled by fluorophore-conjugated streptavidin. Method III is named as Direct Fluorescent Glycan Labeling (DFGL) where target glycans are labeled by direct incorporation of fluorophore-conjugated sialic acid. In all cases, the original terminal sialic acid is removed by neuraminidase treatment. Neu, neuraminidase; STs, sialyltransferases; CMP, cytidine monophosphate; SA, sialic acid. Monosaccharide symbols follow the SNFG (Symbol Nomenclature for Glycans) system (PMID 26543186, Glycobiology 25: 1323–1324, 2015) details at NCBI.

Despite the abundance of N- and O-glycans and their important biological functions, research on these glycans was hampered for the lack of high affinity (Ambrosi, M., Cameron, N.R., et al. 2005) and specific binding reagents (Geisler, C. and Jarvis, D.L. 2011, Sterner, E., Flanagan, N., et al. 2016). The emerge of click chemistry (Kolb, H.C., Finn, M.G., et al. 2001) and metabolic glycan labeling using clickable carbohydrate (Codelli, J.A., Baskin, J.M., et al. 2008, Hsu, T.L., Hanson, S.R., et al. 2007, Kizuka, Y., Funayama, S., et al. 2016) made generic glycan imaging possible. Subsequently, enzymatic glycan labeling using clickable sugars made some specific glycan imaging feasible (Chaubard, J.L., Krishnamurthy, C., et al. 2012, Mbua, N.E., Li, X., et al. 2013, Wu, Z.L., Person, A.D., et al. 2018) (Method I in Fig. 1D). Recently, probing or labeling N-glycans and sT antigen via direct incorporation of biotinylated sialic acids have been reported, respectively, via ST6Gal1 (Capicciotti, C.J., Zong, C., et al. 2017) and ST6GalNAc4 (Wen, L., Liu, D., et al. 2018) (Method II in Fig. 1**D**). Here, we describe the specific imaging of these glycans on HeLa cells via Direct Fluorescent Glycan Labeling (DFGL) where fluorophore-conjugated sialic acids are directly enzymatically incorporated to the target glycans (Method III in Fig. 1**D**).

HeLa cells are derived from human cervical cancer cells (Scherer, W.F., Syverton, J.T., et al. 1953) and are genetically stable (Adey, A., Burton, J.N., et al. 2013). HeLa cells are also known to express CA125, a glycoprotein cancer marker that carries T/sT antigens (Seelenmeyer, C., Wegehingel, S., et al. 2003). By comparing the images of N-glycan and T/sT antigens, we found that these glycans on HeLa cells have different localization, and, more interestingly, the expression of sT antigen on HeLa cells displays random and sporadic patterns that are likely age and stress dependent.

## Results

### Fluorophore Conjugated Sialic Acids can be Directly Introduced to Glycoprotein by Various Sialyltransferases

Fluorophore conjugated CMP-sialic acids (CMP-SA) were synthesized by incubating equivalent CMP-azido-sialic acid (CMP-N_3_-SA) and an alkyne fluorophore via copper (I)-catalyzed azide-alkyne cycloaddition (Rostovtsev, V.V., Green, L.G., et al. 2002). Alexa Fluor^®^ 555 conjugated CMP-sialic acid was first tested on fetal bovine fetuin using various sialyltransferases. The labeling reactions were separated on SDS-PAGE and directly imaged with a regular protein gel imager (Fig. 2A) and a fluorescent gel imager (Fig.2B and 2C). As results, Alexa Fluor® 555 conjugated sialic acid was directly introduced to asialofetuin strongly by ST3Gal1, ST3Gal2 and ST6Gal1, and, weakly by ST6GalNAc1 and ST6GalNAc2. In contrast, the fluorescent sialic acid was directly introduced to fetuin only by ST6GalNAc1, ST6GalNAc2 and ST6GalNAc4. Therefore, ST3Gal1, ST3Gal2, ST6Gal1 showed strict specificity towards asialofetuin and ST6GalNAc4 showed strict specificity towards fetuin, which is consistent to the observation previously achieved with azido-sialic acid (Wu, Z.L., Huang, X., et al. 2016).

**Figure 2.**
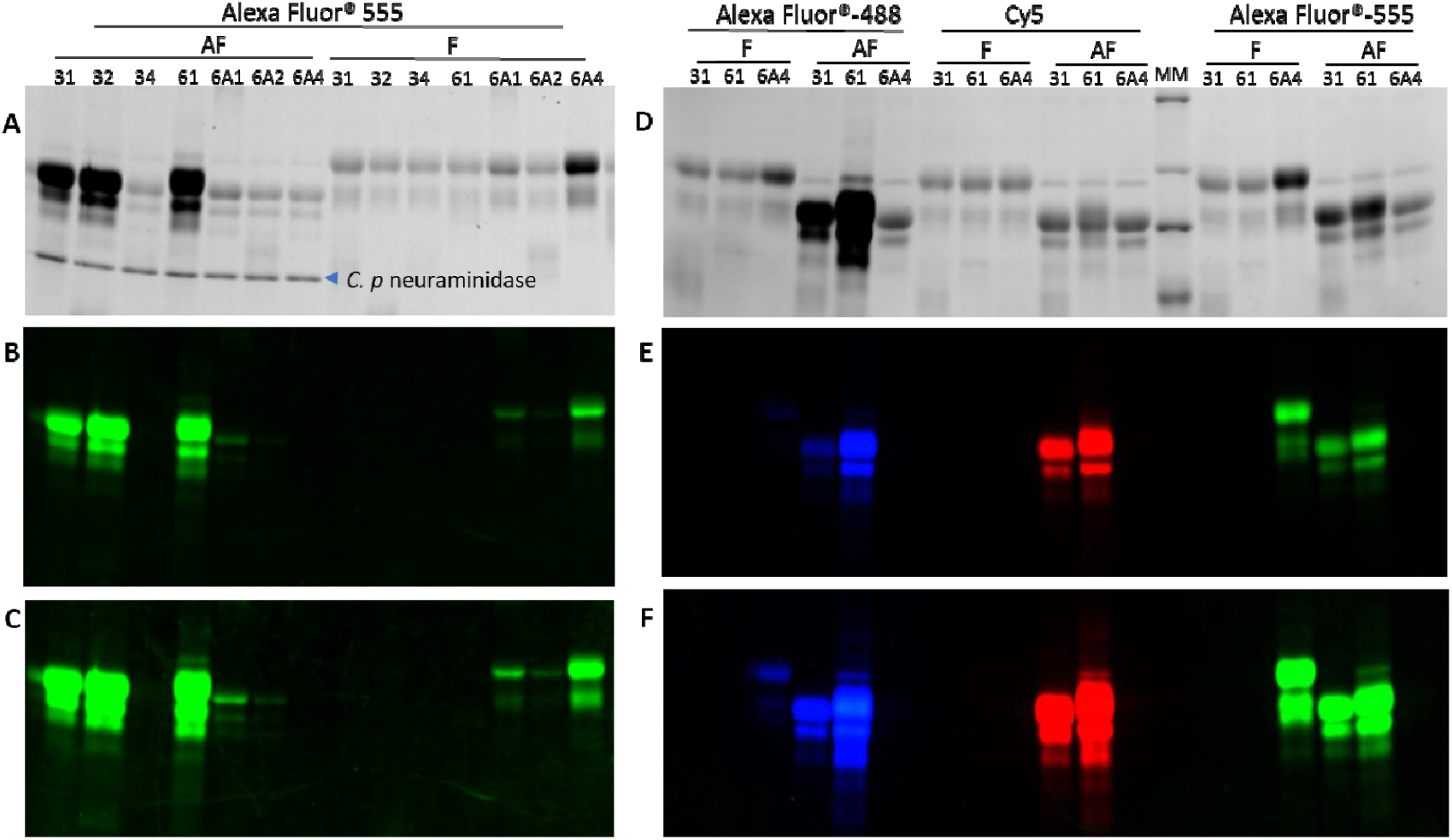
Tolerance of fluorophore conjugated CMP-Sialic acids by various sialyltransferases. Fetal bovine fetuin (F) and asialofetuin (AF) were labeled by various recombinant sialyltransferases with indicated fluorophore conjugated CMP-sialic acids. The labeling reactions were separated on SDS-PAGE and imaged by trichloroethanol (TCE) staining and fluorescent imager. **A, B** and **C** are for a same gel to test the tolerance of Alexa Fluor® 555 by the indicated enzymes. **A** is a TCE image. **B** and **C** are fluorescent images with different contrast. **D E** and **F** are for a same gel to test the tolerance of Alexa Fluor® 488, Alexa Fluor® 555 and Cy5 by the indicated enzymes. **D** is a TCE image. **E** and **F** are fluorescent images with different contrast. 31, ST3Gal1; 32, ST3Gal2; 34, ST3Gal4; 61, ST6Gal1; 6A1, ST6GalNAc1; 6A2, ST6GalNAc2; 6A4, ST6GalNAc4. Same amount of protein (2.5 μg) was loaded into each lane in both **A** and **D**; however, due to the presence of multiple benzene rings in Alexa-Fluor® dyes, Alexa Fluor® 488 and 555 labeled proteins showed significantly increased band intensities in TCE gels. Labeling was success in the presence of *C.P neuraminidase* in panel **A**, suggesting that the neuraminidase has no glycosidase activity on the introduced fluorophore conjugated sialic acids. MM, molecular marker.

ST3Gal1, ST6Gal1 and ST6GalNAc4 were further selected to compare their tolerance on Alexa Fluor® 488, Alexa Fluor® 555 and Cy5 conjugated CMP-SA (Fig. 2D, 2E and 2F). As results, all three dyes were tolerated well by ST3Gal1 and ST6Gal1, but Cy5 was not tolerated by ST6GalNAc4 at all.

### Differential Display of N- and O-glycans on HeLa Cells

For their strict substrate specificities, ST3Gal1, ST6Gal1 and ST6GalNAc4 were then selected to image their substrate glycans on confluent HeLa cells grown in a 96-well plate using Alexa Fluor® 555 conjugated CMP-SA (Fig. 3). Some cells were pretreated with *C. perfringens* neuraminidase (Corfield, A.P., Higa, H., et al. 1983) briefly to remove the existing α2-3 and α2-6 linked sialic acids before imaging. As results, ST3Gal1 and ST6Gal1 produced strong staining only after the neuraminidase treatment, suggesting that Core-1 O-glycans (or T antigen) and N-glycans were largely sialylated on the HeLa cells. In contrast, ST6GalNAc4 only produced strong staining prior to neuraminidase treatment, indicating the presence of sT antigen on the HeLa cells. Interestingly, while ST3Gal1 and ST6GalNAc4 staining was distributed throughout the cells (Fig. 3B and 3C), ST6Gal1 staining was more peripherally localized (Fig. 3. **F**). For comparison, some cells were also imaged by OGT (Fig. 3. **H**), an enzyme that strictly recognizes intracellular compartments, especially nuclei and cytoplasm (Wells, L., Vosseller, K., et al. 2001, Wu, Z.L., Tatge, T.J., et al. 2018). Compared to the intracellular staining by OGT, ST6Gal1 clearly stained the cell surface, which became more obvious when HeLa cells were double imaged by ST6Gal1 and OGT (Supplemental Fig. 1). The layer of glycans on the surface of HeLa cells revealed by ST6Gal1 is comparable to that of the glycocalyx of the endothelial cells revealed by electron microscopy (Ebong, E.E., Macaluso, F.P., et al. 2011). Moreover, in images exhibiting positive staining by ST3Gal1 (Fig. 3. B) and ST6GalNAc4 (Fig. 3.C), some cells were weakly stained or not stained, indicating the variable expression of T/sT antigens by the HeLa cells.

**Figure 3.**
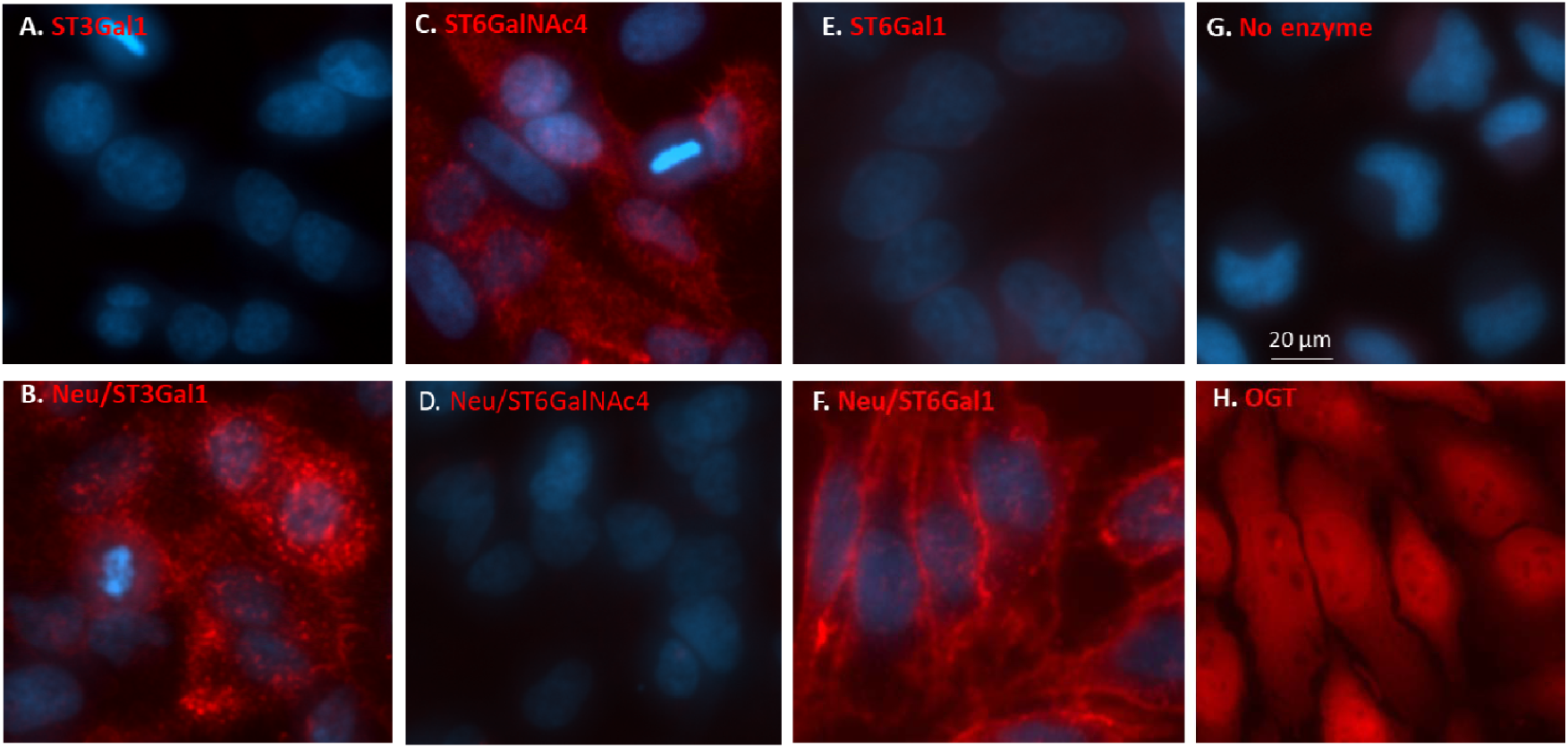
N- and O-glycans exhibit different localization on HeLa Cells. Confluent HeLa cells were imaged by ST3Gal1 (**A** and **B**), ST6GalNAc4 (**C** and **D**), ST6Gal1 (**E** and **F**) before (**A, C, E**) and after (**B, D, F**) *C.p. neuraminidase* treatment. The cells were stained by ST3Gal1 and ST6Gal1 only after *C. perfringens* neuraminidase treatment, suggesting that both Core-1 O-glycan (T antigen) and N-glycan were highly sialylated on these cells. On the contrary, sT antigen was only stained by ST6GalNAc4 prior to *C. p* neuraminidase treatment (**C**). While sT antigen revealed in panels **B** and **C** evenly distributed on cell surface, N-glycans revealed in panel F showed a more peripheral localization. Additionally, not all cells in panel **B** and **C** were equally positive for sT antigen. All images were revealed by Alexa Fluor® 555. For comparison, intracellular contents were stained by OGT in **H** using UDP-azido-GlcNAc that was further reacted to biotin-alkyne and revealed by streptavidin conjugated Alexa Fluor® 555. Nuclei were stained in blue with DAPI.

### Differential Expression of N- and O-glycans Confirmed by Direct Double Imaging

To confirm that O-and N-glycans have different localization, HeLa cells at different confluence were double imaged by ST3Gal1 and ST6Gal1 (Fig. 4), in which Alexa Fluor® 555 conjugated sialic acid was first introduced by ST3Gal1 followed by the introduction of Alexa Fluor® 488 conjugated sialic acid by ST6Gal1. Again, it was observed that N-glycans revealed by ST6Gal1 were preferentially expressed at the cell edges and Core-1 O-glycans were more evenly distributed on cells (also see in Supplemental Fig. 2).

**Figure 4.**
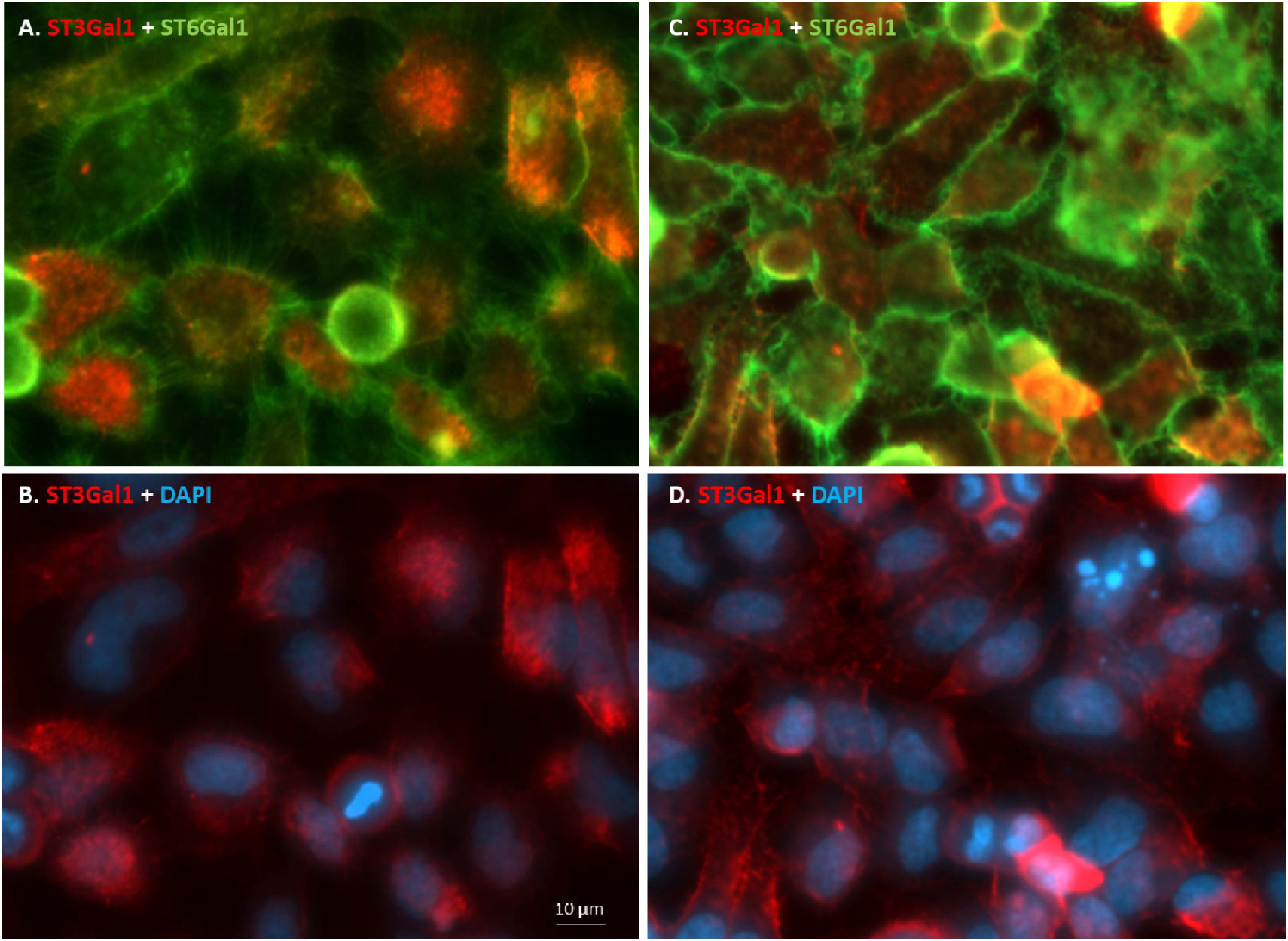
Differential Expression of N-glycans and Core-1 O-glycans Revealed by Double Imaging. HeLa cells at sub-confluence (**A** and **B**) and fully confluence (**C** and **D**) conditions were briefly treated with *C. perfringens* neuraminidase to remove preexisting sialic acids and then imaged for Core-1 O-glycans by ST3Gal1 using Alexa Fluor® 555 conjugated CMP-SA (red) and N-glycans by ST6Gal1 using Alexa Fluor® 488 conjugated CMP-SA (green). **A** and **B** are for the exact same viewing area. **C** and **D** are for the exact same viewing area. Cell nuclei were revealed with DAPI staining in blue in **B** and **D**. N-glycans are expressed preferentially at peripheral positions, especially on protrusions of the cells, compared to Core-1 O-Glycans that are more centralized and evenly distributed on cells.

To better understand the expression of N-glycans, HeLa cells of increasing confluence were imaged by ST6Gal1 (Supplemental Fig. 3). At about 50% culture confluence, N-glycans already displayed preferential distribution at peripheral positions. It seemed that the N-glycans tend to accumulate at the cell edges during growth possibly for cell attachment and migration. This tendency continued until 100% confluence was reached, in which glycocalyx were merged and condensed to ridges between cells (Fig. 4C and Supplemental Fig.3 D).

### Mosaic Expression T/sT antigen and Its Mechanism of Formation

To confirm and better understand the mosaic expression of sT antigen observed in Fig.3C, confluent HeLa cells that were imaged by ST6GalNAc4 were further imaged by OGT to reveal all cells (Fig. 5). Again, sT antigen expression varied significantly from cell to cell, with some cells being strongly positive and some cells being almost completely negative. Occasionally, a positive cell was found to be surrounded by negative cells (circled in Fig. 5B), suggesting that the positive cell be derived from the negative cells, as it is unlikely for a single positive cell being trapped among negative cells during cell division and migration. Mosaic expression of T/sT antigen was also observed on HeLa cells imaged by ST3Gal1 (Supplemental Fig. 4).

**Figure 5.**
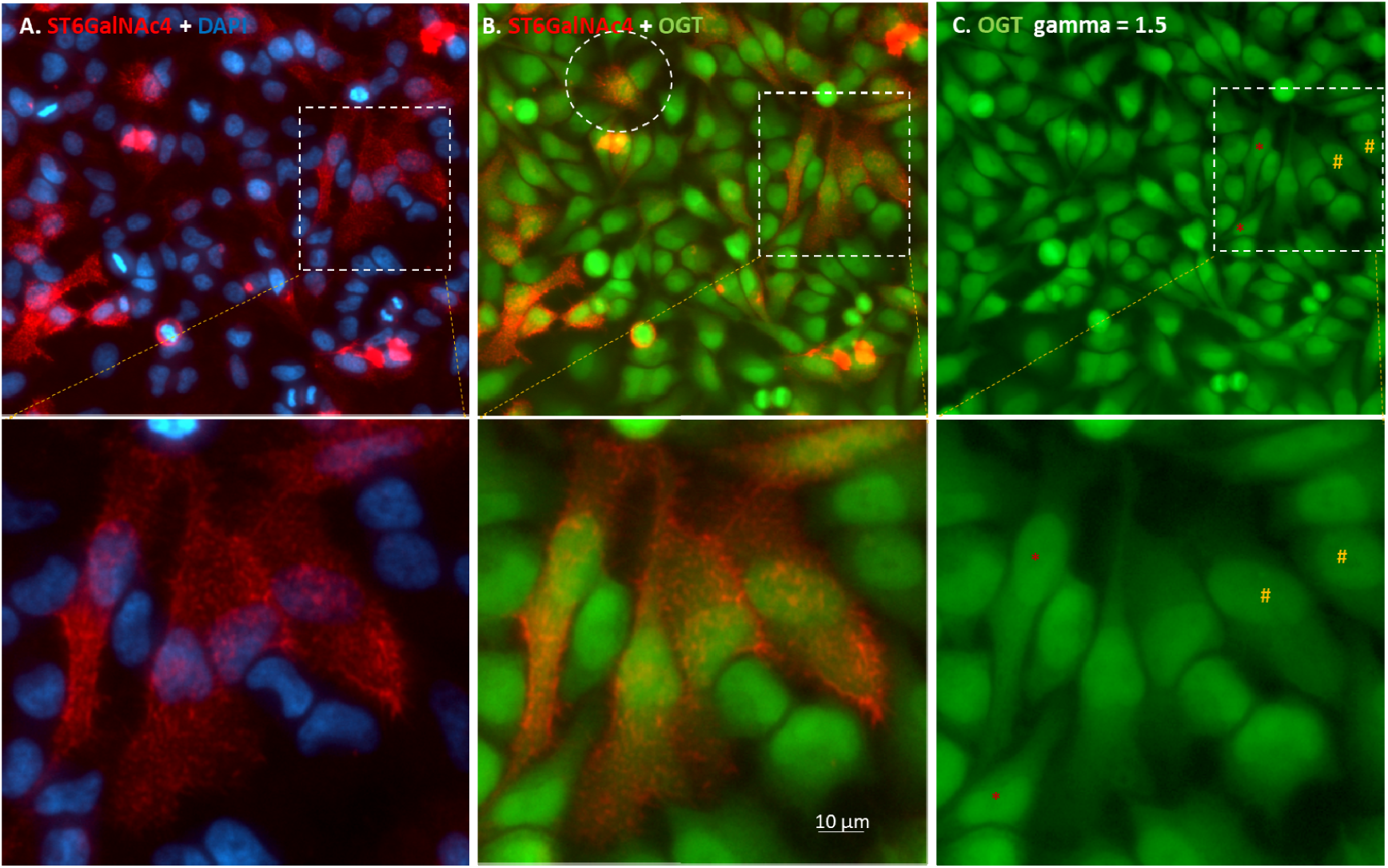
Mosaic sT Antigen Expression on HeLa cells. Confluent HeLa cells were imaged for sT antigen by ST6GalNAc4 (red, via Alexa fluor® 555). Nuclei were revealed in blue with DAPI. Intracellular contents were revealed by OGT with UDP-azido-GlcNAc and Alexa Fluor® 488 via click chemistry (green). Panel **A, B** and **C** are from a same viewing area. A single positive cell surrounded by negative cells is circled in panel **B**. Cells marked with * and # in panel **C** are for neighboring cells that exhibit striking different intensities on sT antigen expression in panel **B**.

HeLa cells are cancerous cells. The expression of sT carbohydrate cancer antigen on HeLa cells is not surprising (Seelenmeyer, C., Wegehingel, S., et al. 2003). However, the mosaic expression of sT antigen is intriguing. Two hypotheses were postulated to explain this phenomenon. The first is that positive cells were derived from negative cells, which is suggested by Fig. 5. The second is that the initial HeLa cell population were heterogeneous and contained positive subpopulation. To test the second hypothesis, HeLa cells from the above experiments were subject to further clonal selection, by which single cells were selected and cultivated under the same conditions for 11-14 days to form individual colonies. When imaged by ST6GalNAc4 at day 11, sT antigen positive cells were sporadically found throughout the colonies but more abundantly around the centers of these colonies (Fig. 6). More sT antigen positive cells were found around the centers of these colonies at day 14 (Supplemental Fig. 5). Since cells around the centers of the colonies are older and more crowded (more stress) than cells at peripheral positions, the above observations together argue that sT antigen positive cells were derived from sT antigen negative cells and sT antigen expression is age/stress dependent.

**Figure 6.**
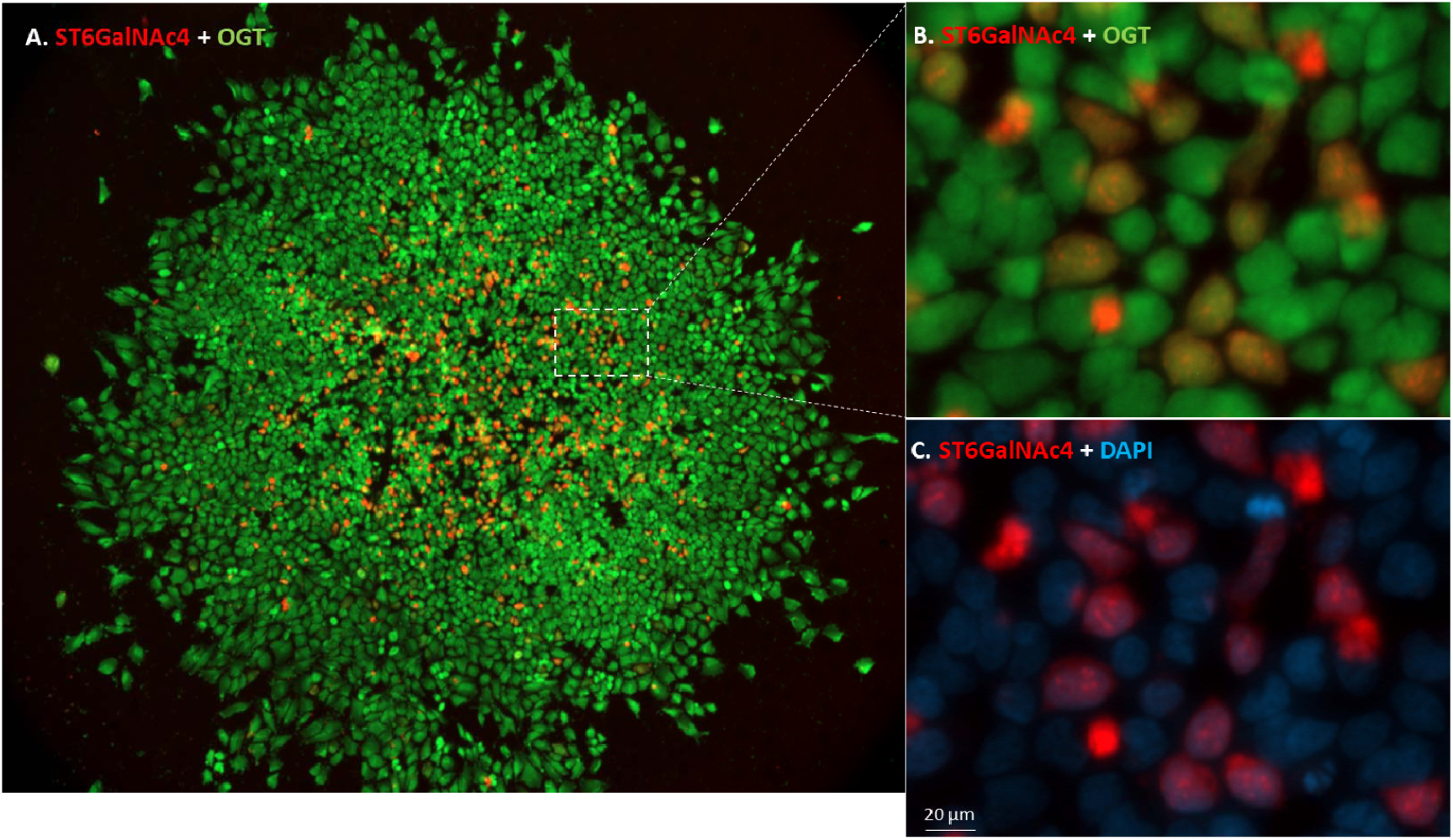
Expression of sT antigen on a colony of HeLa cells that were derived from a single cell. A HeLa cell population was plated for single cell isolation and grown in DMEM medium (with 10% FBS) for 11 days at 37 °C with 5% of CO_2_ until a single colony formed in the well. The colony was imaged by ST6GalNAc4 with Alexa Fluor® 555 conjugated CMP-SA in red and countered imaged by OGT with UDP-azido-GlcNAc and Alexa Fluor® 488 in green. Nuclei were stained with DAPI in blue. The sT antigen positive cells are more abundant at the center compared to the edges of the colony. As the cells at the center of the colony were older and more crowded than the cells at the edges of the colony, this result may suggest that the expression of sT antigen is age and stress dependent. **A**, the double staining of a single colony. **B**, the magnified region indicated in **A**. **C**, the magnified region in **A** showing ST6GalNAc4 and DAPI staining.

## Discussion

In this report, we describe the specific imaging of N- and O-glycans on HeLa cells and present evidence that these glycans have distinct expression patterns. While O-glycans such as T/sT antigens are more evenly distributed on cells, N-glycans tend to be localized at peripheral positions. While N-glycans are expressed on all cells, T/sT antigens expression is sporadic and likely age/stress dependent.

The distinct localizations of N- and O-glycans suggest that they are differentially conjugated to glycoproteins on cell surface, and individual glycoproteins may be enriched with one type of glycan but not the other. The distinct localizations also suggest that these glycans may have different functions. For instance, the N-glycans expressed at cell edges could be involved in cell-cell and cell-ECM interactions to facilitate cell adhesion and migration, and the O-glycans displayed on cell surface could be more involved in ligand-receptor interaction and self-protection against pathogens. Consistently, integrins that are extensively modified with N-glycans are known to be localized at cell edges and to mediate cell-cell and cell-matrix interactions (Hynes, R.O. 2002), and, mucins that are heavily O-glycosylated are usually expressed on epithelial cell surface where they fulfill a protective role by binding to pathogens (Hattrup, C.L. and Gendler, S.J. 2008, Linden, S.K., Sheng, Y.H., et al. 2009).

As a technique, DFGL has several advantages over previous methods for glycan labeling/imaging. First, the method is more convenient and time-saving by involving only a single step of enzymatic reaction prior to cell imaging and allowing direct imaging of an SDS gel without the time-consuming membrane transfer step in Western blotting. Second, this method has eliminated the side effect of oxidative cleavage of target proteins and is more suitable for live cell imaging by avoiding the usage of copper ion of traditional click chemistry reaction. Third, this method allows double or multiple imaging of different glycans simultaneously without cross-staining of these glycans by avoiding the common intermediate step of biotin-streptavidin reaction. Fourth, since fluorophore is introduced only by enzymatic reaction, the labeling/detection of a certain glycan is more specific and has lower background. In summary, DFGL is a convenient tool for specific glycan detection, labeling and imaging. It is also foreseeable that this technique can be applied to detect other types of glycans using different glycosyltransferases and fluorophore-conjugated sugars as donor substrates.

The fact that specific glycan structures can be reproducibly generated by normal cells without templates suggests that the Golgi apparatus has an extremely well-organized structure with the synthetic glycosyltransferases localized in precise order. It is hypothesized that this ordered structure in HeLa cells or pre-cancerous cells is rather fragile and easy to be disrupted under the effects of age and/or stress, leading to premature synthesis of abnormal shortened glycan epitopes.

Cancers were considered as genetic diseases (Alexandrov, L.B., Nik-Zainal, S., et al. 2013). However, recent work suggests that mutations of oncogenes arise in normal tissues may not necessarily lead to cancer (Martincorena, I., Fowler, J.C., et al. 2018). Direct causes of cancerous phenotype may need additional factors, such as environmental triggers (Chanock, S.J. 2018). Conversely, a cancerous phenotype may not be a direct outcome of genetic mutations. Accordingly, our results on sT antigen imaging strongly argue that its expression is not due to immediate genetic mutation but caused by disrupted posttranslational machinery. Recently, glycan cancer antigens have been targeted for cancer immunotherapy (Posey, A.D., Jr., Schwab, R.D., et al. 2016, Steentoft, C., Migliorini, D., et al. 2018). A way for imaging or detecting these glycan cancer antigens may help us to better design therapeutic strategies.

## Material and Methods

CMP-Azido-Sialic acid, UDP-Azido-GlcNAc, Biotinylated Alkyne, DAPI, recombinant human ST3Gal1, ST3Gal2, ST3Gal4, ST6Gal1, ST6GalNAc1, ST6GalNAc4, OGT and *C. perfringens* Neuraminidase were from Bio-Techne. Alkyne-Alexa Fluor® 488, alkyne-Alexa Fluor®555, streptavidin-Alexa^®^ Fluor 555 were from Thermo Fisher Scientific. Cy5-alkyne, ascorbic acid, fetal bovine fetuin and asialofetuin and all other chemical reagents were from Sigma-Aldrich.

### Preparation of fluorescent conjugated CMP-sialic acid

Fluorophore conjugated CMP-sialic acids were prepared by incubating equivalent CMP-Azido-Sialic acid (CMP-N_3_-SA) and an alkyne-conjugated fluorophore via copper (I)-catalyzed azide-alkyne cycloaddition. As a typical reaction, 5 mM of CMP-N_3_-SA was mixed with 5 mM of Cy5-alkyne in the presence of 0.1 mM of Cu^2+^ and 1 mM of ascorbic acid. The reaction was kept at room temperature for 2 hours. The fluorophore conjugated CMP-sialic acids were then purified on a HiTrap® Q HP (GE Healthcare) column and eluted with a 0-100% gradient of NaCl elution buffer (150 NaCl, 25 mM Tris at pH 7.5). The fluorophore conjugated CMP-SA was collected based on color exhibition and UV absorption as the conjugated CMP-SA was vivid in color and had UV absorption at 260 nm. Alexa Fluor® 555 conjugated CMP-SA, Alexa Fluor® 488 conjugated CMP-SA, Cy5 conjugated CMP-SA were prepared and purified accordingly and finally concentrated to >0.1 mM by a speed-vacuum concentrator.

### Protein labeling and imaging with fluorophore conjugated CMP-sialic acid

For a typical glycoprotein labeling reaction, 1 to 5 μg target protein was mixed with 0.2 nmol fluorophore conjugated CMP-SA and 0.2 μg of a sialyltransferase in 30 μL buffer of 25 mM Tris pH7.5, 10 mM MnCl_2_. The mixture was incubated at 37°C for 20 minutes. In the case that the preexisting sialic acid of a glycoprotein needed to be removed, 0.01 μg of *C. perfringens* Neuraminidase was also added into the reaction. The neuraminidase had no activity on fluorophore conjugated sialic acid and was not necessary to be removed after the reaction. The reaction was then separated by sodium dodecyl sulfate–polyacrylamide gel electrophoresis (SDS-PAGE) and the gel was directly imaged using a fluorescent imager FluorChem M (ProteinSimple, Bio-techne).

### Cell Culture

HeLa cells (ATCC # CCL2) were grown in MEM NEAA Earle’s Salts (Irvine Scientific, #9130), supplemented with 10% fetal bovine serum (Corning, #35-015-CV), 2 mM L-glutamine, 100 units/mL penicillin and 0.1 mg/mL streptomycin (Sigma-Aldrich, #G6784) in an incubator at 37°C and with 5% CO_2_. Upon confluence, cells were trypsinized and seeded in a 96-well cell culture plate and grown to a desired confluence.

### Cell imaging with Direct Fluorescent Glycan Labeling (DFGL)

For a typical cell imaging on a 96-well plate, the medium in each well was removed and replaced with a labeling mixture containing 0.2 nmol of a fluorophore-conjugated CMP-SA, 0.5 µg of a sialyltransferase, 30 µL labeling buffer (25 mM Tris pH 7.5, 150 mM NaCl, 10 mM Mn^2+^) and 20 µL of the original medium, and the cells were then incubated at 37°C for 20 minutes. For cell imaging using ST3Gal1 or ST6Gal1, cells in each well were pretreated with 1 µg of *C.p* Neuraminidase directly in the medium at 37°C for 10 minutes prior to the labeling. After the labeling, the cells were washed three times quickly each with 150 µL 25 mM Tris pH 7.5, 150 mM NaCl. For double imaging with another fluorophore, the cells were incubated with a second labeling mixture at 37°C for 20 minutes and then washed thoroughly, followed by fixation in 4% paraformaldehyde for 10 minutes at room temperature. The fixed cells were then stored in the buffer of 25 mM Tris pH 7.5, 150 mM NaCl for imaging.

### Intracellular content imaging with OGT

After fixation, for labeling the intracellular contents by OGT, a labeling mixture of 1 nmol of UDP-Azido-GlcNAc, 1 µg of OGT, 50 µL of the labeling buffer (25 mM MES, 0.5% (w/v) Triton® X-100, 2.5 mM MgCl_2_, 10 mM MnCl_2_, 1.25 mM CaCl_2_ and 0.75 mg/mL of BSA at pH 7.0) was applied to each well of a 96-well plate. Triton® X-100 was added into the reaction to permeabilize the cell membrane. The reaction was kept at 37°C for 60 minutes. After the labeling mixture was removed completely, the cells were washed three times in a washing buffer (25 mM Tris pH 7.5, 150 mM NaCl). The labeled cells were then subject to click chemistry reaction by incubating the cells in 0.1 mM Cu^2+^, 1 mM ascorbic acid and 0.1 mM Biotinylated Alkyne at room temperature for 30 minutes. After the click reaction was removed completely, the cells were incubated in 50 μL of 10 μg/mL streptavidin-Alexa^®^ 555 or 488 in the washing buffer for 15 minutes. DAPI (10 μM) was added at this stage to stain the cell nuclei. The cells were then washed thoroughly and stored in the washing buffer for imaging.

### Equipment and Settings

All cell images were captured on an AXIO Observer microscope (ZEISS) with a ZEISS Axiocam 506 mono camera and Zen 2 Pro software. Images were captured simultaneously through different fluorescent channels. Exposure time for each channel ranged from 100 ms to 4000 ms. For best viewing, obtained images from each channel was usually normalized to its highest pixel value using the best-fit option of the software. In the case that gamma rating was changed, it was indicated in figure legends. For taking fluorescent image of an SDS gels using FluorChem M, multiple fluorescent channel RGB or single fluorescent channel was selected based on the incorporated fluorescent dyes, and the exposure time was set at Auto, and it was indicated in the figure legends when contrast was manually enhanced.

## Acknowledgments

We want to acknowledge numerous colleagues who have made contributions to this manuscript through product development, quality control and assurance, and product support.

