## supplemental figures 1-5 for "Differential Expression of N- and O-glycans on HeLa Cells-Revealed by Direct Fluorescent Glycan Labeling with Recombinant Sialyltransferases"

### Slide 1
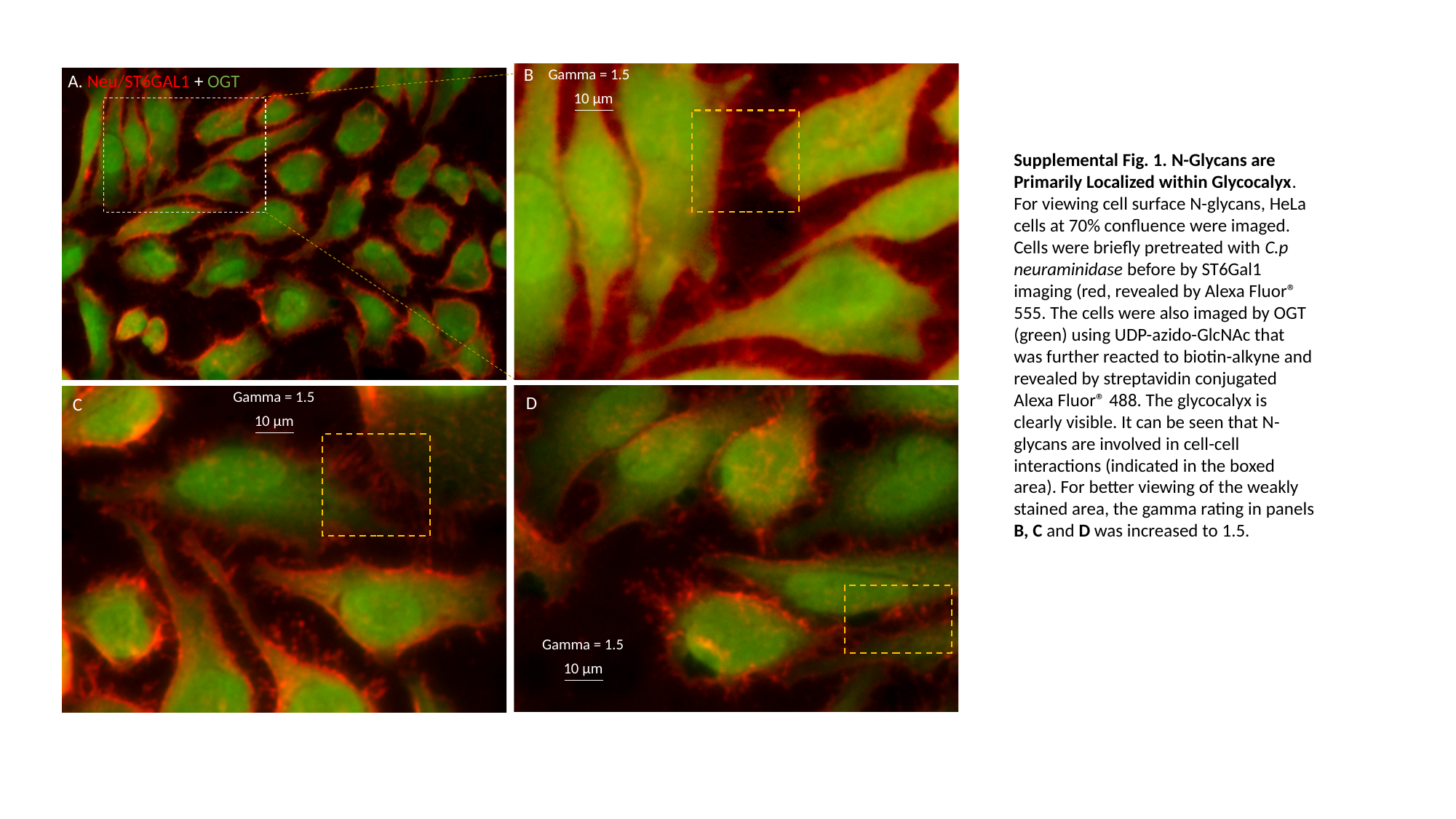

B
Gamma = 1.5
A. Neu/ST6GAL1 + OGT
10 µm
Supplemental Fig. 1. N-Glycans are Primarily Localized within Glycocalyx. For viewing cell surface N-glycans, HeLa cells at 70% confluence were imaged. Cells were briefly pretreated with C.p neuraminidase before by ST6Gal1 imaging (red, revealed by Alexa Fluor® 555. The cells were also imaged by OGT (green) using UDP-azido-GlcNAc that was further reacted to biotin-alkyne and revealed by streptavidin conjugated Alexa Fluor® 488. The glycocalyx is clearly visible. It can be seen that N-glycans are involved in cell-cell interactions (indicated in the boxed area). For better viewing of the weakly stained area, the gamma rating in panels B, C and D was increased to 1.5.
Gamma = 1.5
D
C
10 µm
Gamma = 1.5
10 µm

### Slide 2
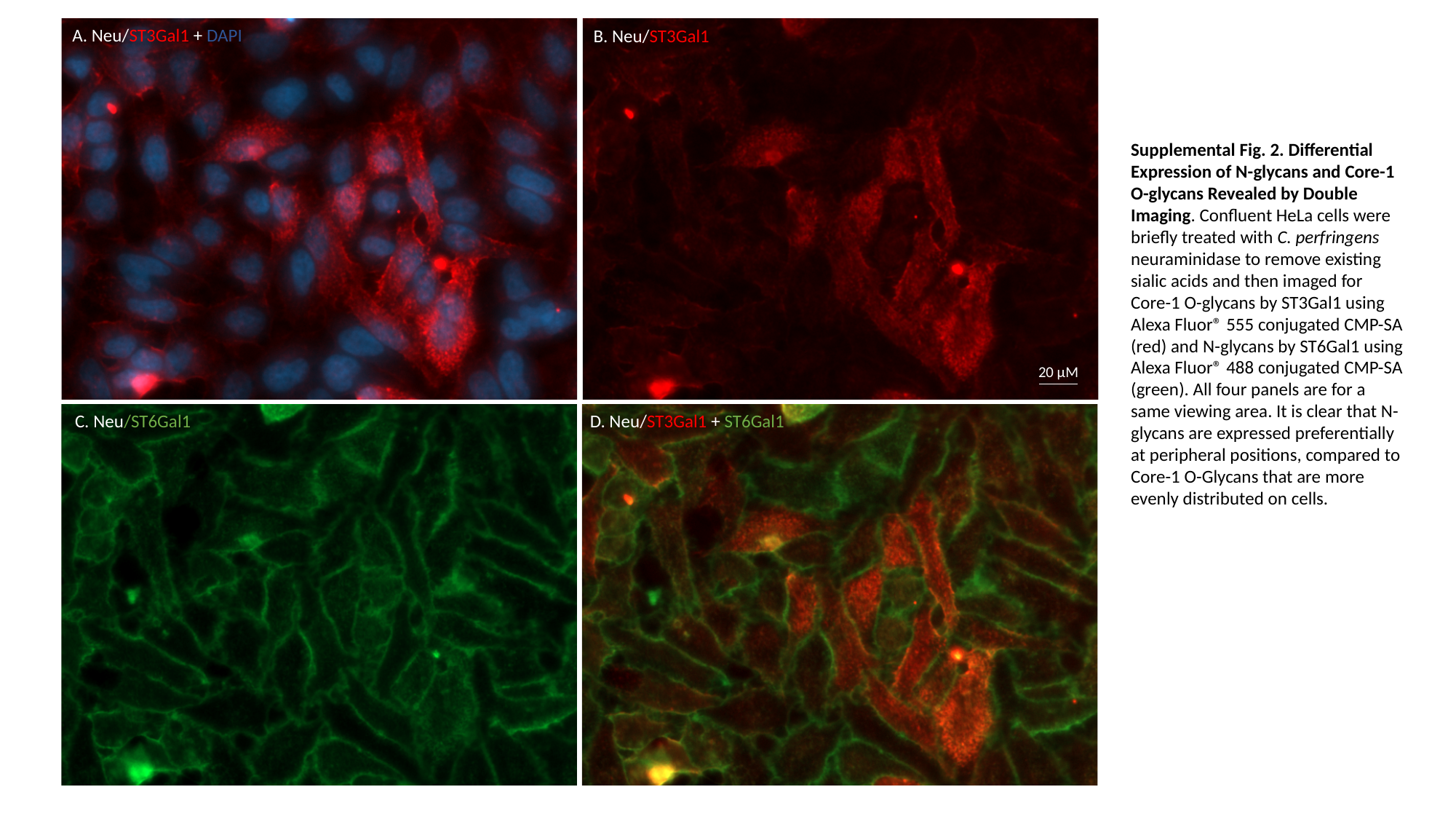

A. Neu/ST3Gal1 + DAPI
B. Neu/ST3Gal1
Supplemental Fig. 2. Differential Expression of N-glycans and Core-1 O-glycans Revealed by Double Imaging. Confluent HeLa cells were briefly treated with C. perfringens neuraminidase to remove existing sialic acids and then imaged for Core-1 O-glycans by ST3Gal1 using Alexa Fluor® 555 conjugated CMP-SA (red) and N-glycans by ST6Gal1 using Alexa Fluor® 488 conjugated CMP-SA (green). All four panels are for a same viewing area. It is clear that N-glycans are expressed preferentially at peripheral positions, compared to Core-1 O-Glycans that are more evenly distributed on cells.
20 µM
C. Neu/ST6Gal1
D. Neu/ST3Gal1 + ST6Gal1

### Slide 3
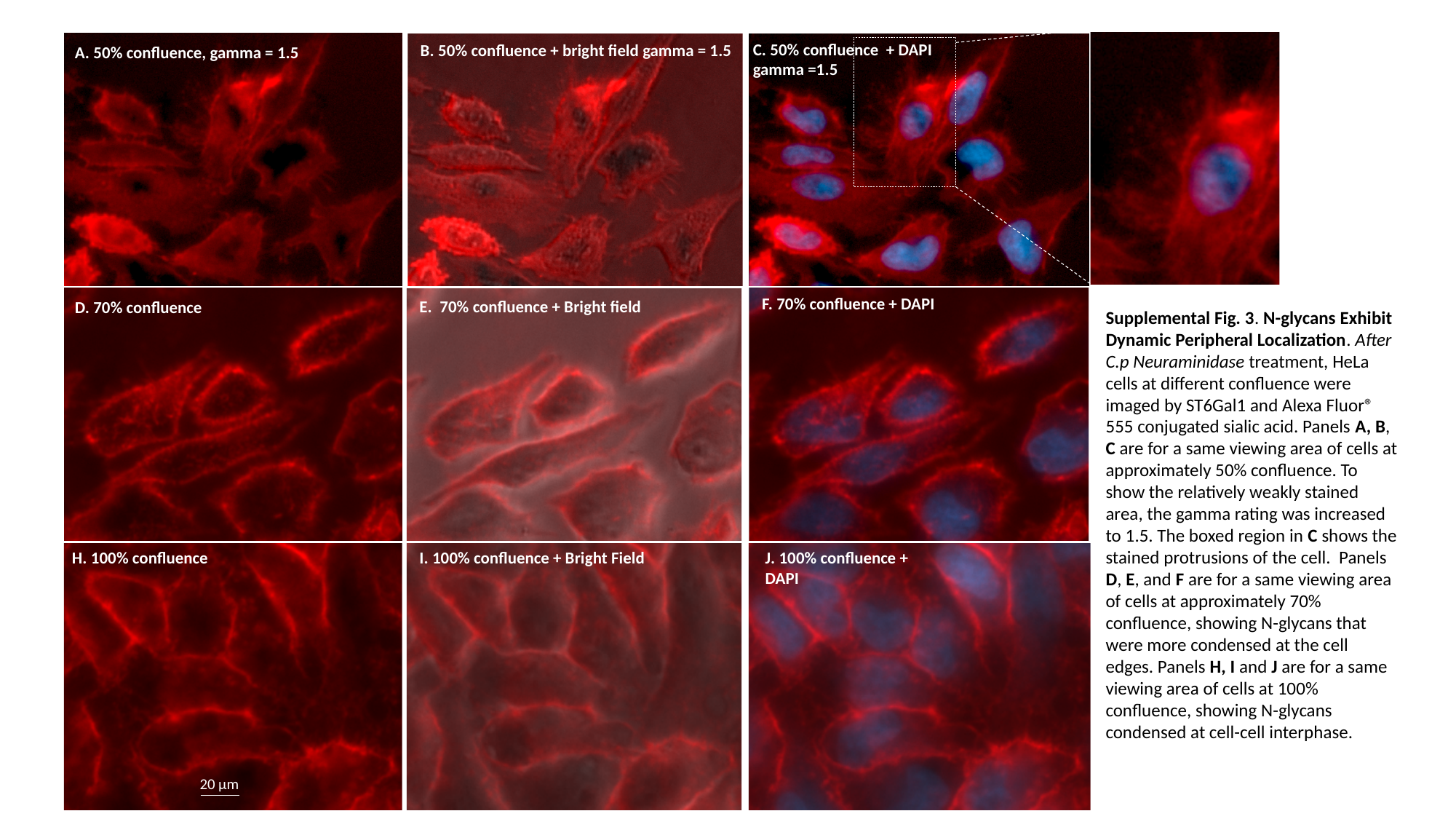

C. 50% confluence + DAPI gamma =1.5
B. 50% confluence + bright field gamma = 1.5
A. 50% confluence, gamma = 1.5
F. 70% confluence + DAPI
E. 70% confluence + Bright field
D. 70% confluence
Supplemental Fig. 3. N-glycans Exhibit Dynamic Peripheral Localization. After C.p Neuraminidase treatment, HeLa cells at different confluence were imaged by ST6Gal1 and Alexa Fluor® 555 conjugated sialic acid. Panels A, B, C are for a same viewing area of cells at approximately 50% confluence. To show the relatively weakly stained area, the gamma rating was increased to 1.5. The boxed region in C shows the stained protrusions of the cell. Panels D, E, and F are for a same viewing area of cells at approximately 70% confluence, showing N-glycans that were more condensed at the cell edges. Panels H, I and J are for a same viewing area of cells at 100% confluence, showing N-glycans condensed at cell-cell interphase.
ST6Gal1
H. 100% confluence
I. 100% confluence + Bright Field
J. 100% confluence + DAPI
20 µm

### Slide 4
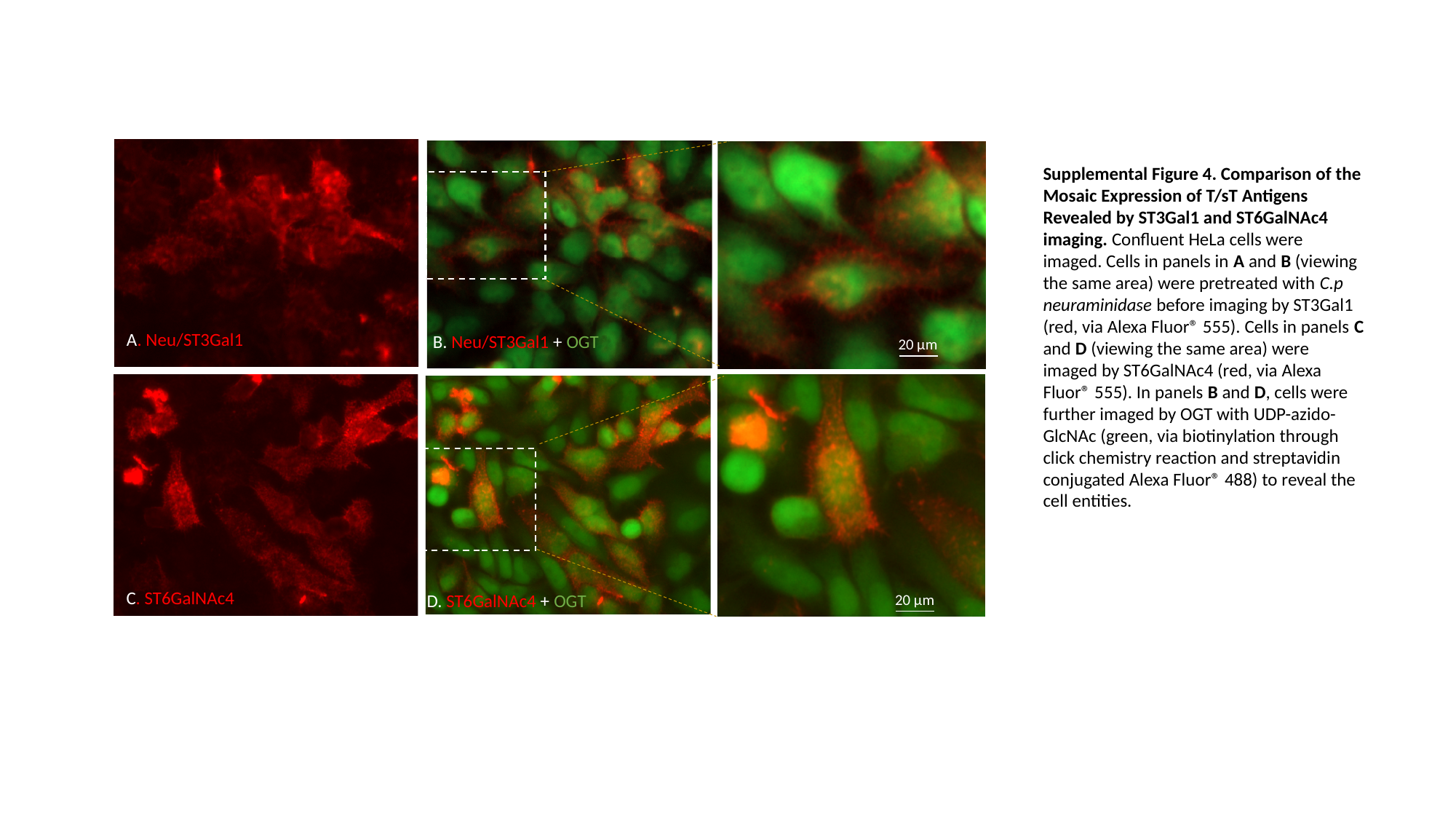

Supplemental Figure 4. Comparison of the Mosaic Expression of T/sT Antigens Revealed by ST3Gal1 and ST6GalNAc4 imaging. Confluent HeLa cells were imaged. Cells in panels in A and B (viewing the same area) were pretreated with C.p neuraminidase before imaging by ST3Gal1 (red, via Alexa Fluor® 555). Cells in panels C and D (viewing the same area) were imaged by ST6GalNAc4 (red, via Alexa Fluor® 555). In panels B and D, cells were further imaged by OGT with UDP-azido-GlcNAc (green, via biotinylation through click chemistry reaction and streptavidin conjugated Alexa Fluor® 488) to reveal the cell entities.
A. Neu/ST3Gal1
B. Neu/ST3Gal1 + OGT
20 µm
C. ST6GalNAc4
D. ST6GalNAc4 + OGT
20 µm

### Slide 5
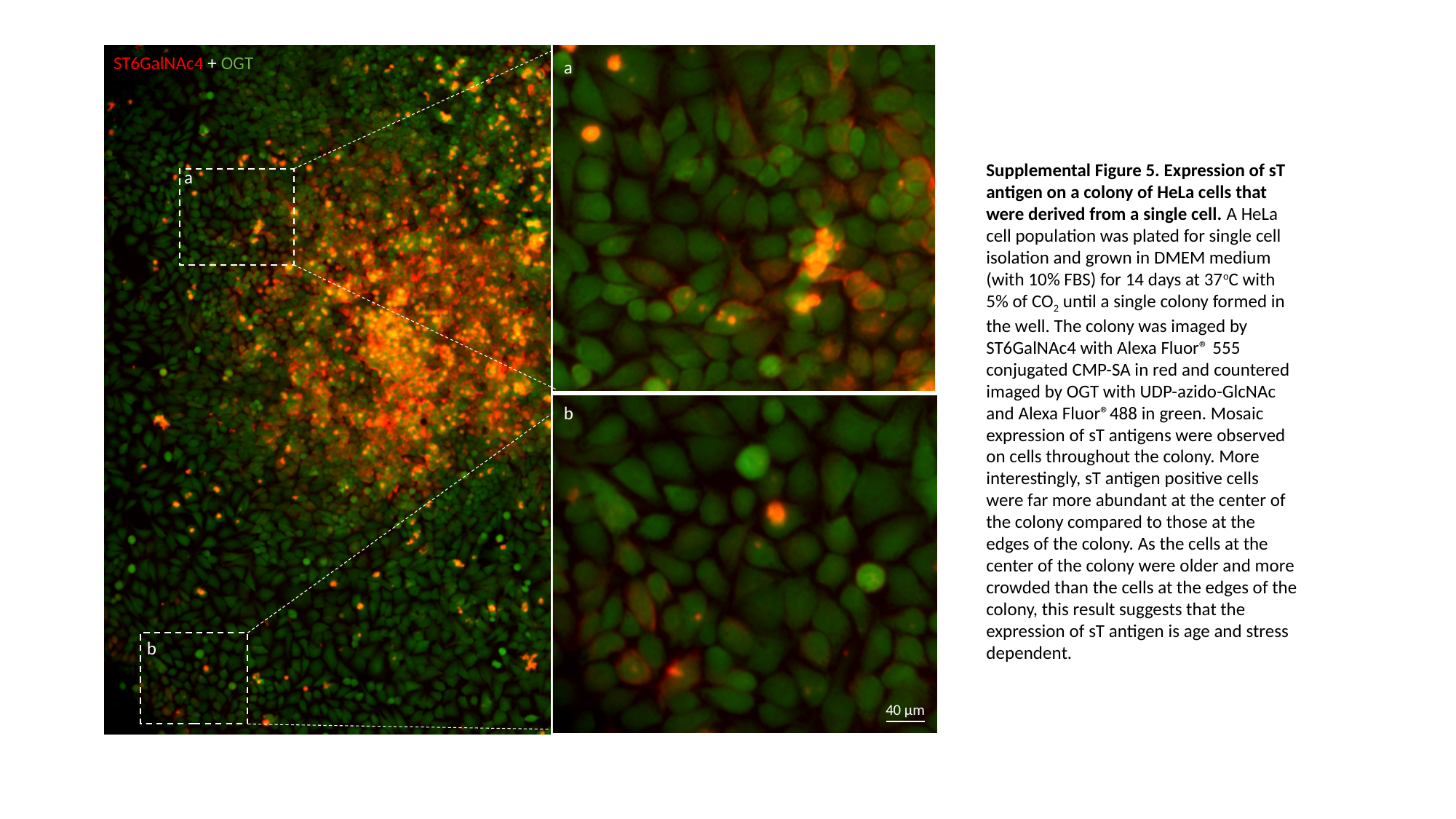

ST6GalNAc4 + OGT
a
Supplemental Figure 5. Expression of sT antigen on a colony of HeLa cells that were derived from a single cell. A HeLa cell population was plated for single cell isolation and grown in DMEM medium (with 10% FBS) for 14 days at 37^o^C with 5% of CO_2_ until a single colony formed in the well. The colony was imaged by ST6GalNAc4 with Alexa Fluor® 555 conjugated CMP-SA in red and countered imaged by OGT with UDP-azido-GlcNAc and Alexa Fluor®488 in green. Mosaic expression of sT antigens were observed on cells throughout the colony. More interestingly, sT antigen positive cells were far more abundant at the center of the colony compared to those at the edges of the colony. As the cells at the center of the colony were older and more crowded than the cells at the edges of the colony, this result suggests that the expression of sT antigen is age and stress dependent.
a
b
b
40 µm
